# Precise 3D Tracking of Highly Non-planar Eukaryotic Flagellar Beating Patterns using Digital Holographic Microscopy

**DOI:** 10.1101/2025.07.28.666810

**Authors:** Patryk Nienaltowski, Jonasz Slomka, Federica Miano, Thomas Kiørboe, Clara Martínez-Pérez, Tristan Colomb, Yves Emery, Roman Stocker

## Abstract

Precise tracking of the rapid and complex three-dimensional movement of eukaryotic flagella is important for understanding their roles in cellular motility, sensory functions, and resource acquisition. Yet, achieving accurate 3D kinematic reconstruction of flagellar beating patterns, particularly highly non-planar ones, remains challenging. Here we present holoV3C, a method based on Digital Holographic Microscopy (DHM) that allows precise, label-free 3D tracking of highly non-planar eukaryotic flagella with high temporal resolution. This algorithm leverages phase anomaly detection to provide a combination of high temporal and axial resolution, with 0.25 μm for beating mouse sperm flagella and down to 53 nm for polystyrene particles, across large sampling volumes in a computationally efficient manner. Algorithmic validation is performed by tracking mouse sperm flagella over time, capturing approximately 600 points along a single flagellum to achieve high axial resolution. Furthermore, we apply holoV3C to reconstruct the highly non-planar beating dynamics of the 200-nm-diameter flagellum of the protist *Reclinomonas americana* with a temporal resolution of 200 frames per second. By enabling 3D tracking of non-planar eukaryotic flagella, holoV3C can yield important insights to advance our understanding of flagellar dynamics, opening new avenues in the study of microorganism motility and its ecological roles.

## 1. Introduction

Movement is the basis of life [1], including the life of microorganisms in aquatic environments where motility is critical for survival [2, 3]. Flagella play a central role by facilitating behaviors such as resource acquisition and evasion from predators [4]. They are found in nearly all eukaryotic lineages, from unicellular organisms to the gametes of most multicellular animals, including mammals [5]. Understanding flagellar movement and how it translates into cell motility is key to understanding the interface between processes at the level of cellular signaling and biophysics and the wider functions of cells at the level of behavior and ecology. For instance, the intrinsic curvature or chiral beating dynamics of flagella can play a role in cell navigation through rheotactic steering [6]. Tracking flagellar beating can also shed light on how human sperm respond to stimuli, providing insights into reproductive biology [7]. Moreover, flagella play a key role not only in escape from predation, but also in the capture of microorganisms by predators [4].

Eukaryotic flagella are complex structures that often extend several micrometers in length while having a diameter as small as 200 nanometers. They can oscillate at frequencies up to 100 Hz [8, 9], underscoring the challenges in measuring and tracking their three-dimensional (3D) shape. Furthermore, the tracking process is complicated not only by the flagellum’s minuscule size, near the optical limit, but also by its often low optical contrast. The need for high temporal resolution to capture the rapid oscillations of flagella imposes stringent demands on optical systems and necessitates the development of tools capable of accurately characterizing their dynamic 3D beating patterns across varying structures, dimensions, and behaviors. Despite the importance of quantifying flagellar motion, existing imaging methods often fall short of these requirements, as they struggle to balance spatial precision, temporal resolution, volumetric sampling, and computational efficiency.

Much of the existing research on eukaryotic flagella movement relies on two-dimensional (2D) observations, capturing only the projections of inherently 3D motility, or is based on analysis of only a narrow portion of the sample restricted by the microscope’s depth of field [10]. This reliance on 2D projections leads to a loss of information, potentially distorting the interpretation and quantitative analysis of flagellar motion. For example, complex helical movements may appear linear or planar when viewed in 2D. Additionally, restricting measurements to a subset limited by the microscope’s depth of field impairs the study of flagella with vertical components to their motion, thereby excluding the analysis of the 3D beating patterns exhibited by many eukaryotic cells. These 2D measurements cannot capture the intricate dynamics of flagellar movement, potentially overlooking critical aspects of microbial motility. Accurate characterization of natural flagellar motion, with all its spatial intricacies, thus necessitates the use of 3D tracking techniques.

The need to address these challenges has led to the development of various 3D methodologies. Although previous efforts have provided valuable insights, their applicability has often been constrained by methodological limitations. The challenges arise not only from limitations in imaging techniques but also from the computational complexity of processing the resulting data. Existing 3D imaging methods, such as light-sheet and confocal microscopy, offer precise 3D measurements but require sample labeling and are constrained by limited temporal resolution and small sampling volumes. Similarly, previous methods for capturing 3D flagellar beating, including multifocal microscopy [11–13], defocused 2D microscopy [6, 14], and light microscopy with piezo-driven objectives [15, 16], have demonstrated some effectiveness but have intrinsic constraints, which may include limited sampling volume, the requirement for labeling, the need for extensive calibration, or reliance on physical scanning to construct 3D stacks, which limits temporal resolution and impacts measurement accuracy. Collectively, these limitations make tracking flagellar beating more challenging and less accurate, hindering our ability to precisely measure the dynamic 3D beating patterns of eukaryotic flagella.

There is thus a need for a method that integrates high spatiotemporal resolution with large sampling volumes and computational efficiency to accurately capture the rapid, 3D movements of eukaryotic flagella. Digital holographic microscopy (DHM) [17] represents a promising solution by enabling rapid volumetric and digital refocusing through sample depths of millimeters [18]. Previous DHM applications for 3D tracking have primarily focused on particle tracking [19–22] and, in the context of sperm cell motility, on reconstructing the 3D trajectories of sperm heads, while the analysis of flagellar dynamics has been relatively underexplored [23–25]. Recent advances have demonstrated the potential of DHM for reconstructing 3D flagellar beating, representing a significant step forward in the characterization of flagellar dynamics [26–30]. However, while most experimental studies have largely concentrated on the quasi-planar beating patterns of sperm flagella, a gap remains for methodologies capable of precisely capturing the highly non-planar, 3D dynamics characteristic of many flagellar waveforms, such as those found in phagotrophic flagellates crucial to marine food webs [4]. In this context, the existing approaches face several limitations, including high computational complexity, restricted sample volumes, and suboptimal axial resolution.

In this work, we present a method for 3D tracking of eukaryotic flagella using holographic imaging. Our approach integrates high spatiotemporal resolution with extensive sampling volumes while eliminating the need for sample labeling. We experimentally validate our method by measuring 3D beating kinematics of flagella with distinct morphological characteristics and motility patterns, namely mouse sperm cells, known for their quasi-planar beating, and the protist *Reclinomonas americana* (Discoba, Jakobida), which exhibits highly non-planar flagellar waveforms. Our results highlight the method’s capacity for precise determination of 3D positions (*x,y,z*) for individual points along the flagellum, and illustrate the computational efficiency of our approach. The combination of high spatiotemporal resolution with large sampling volume and operational efficiency makes this method a valuable tool for the precise tracking of 3D flagellar kinematics across diverse biological contexts.

## 2. Results

### 2.1. 3D tracking method

This paper introduces a 3D tracking method called holoV3C, specifically designed for the precise tracking of eukaryotic cells and their microstructures such as flagella. The name “holoV3C” reflects its fundamental functionality: the application of holographic (holo) techniques to extract the three-dimensional (3) position of a cell by analyzing one-dimensional amplitude vectors (VEC) along the optical axis. The inclusion of “3” within “VEC” underscores the method’s ability to achieve precise single-point (pixel-level) localization in 3D space.

The method is structured into four sequential core modules: acquisition, lateral localization, axial localization, and interpretation (Figure 1). The acquisition module employs digital holographic microscopy to capture holograms of the sample volume, creating a 3D record of the specimen that is then subjected to holographic reconstruction [17]. The second module, lateral localization, extracts the (*x,y*) positions of points along the flagellum using phase *z*-stack and image processing techniques. The third module is axial localization, which determines the *z* positions along the optical axis for the lateral (*x,y*) coordinates found in the previous module. The second and third modules are interlinked, they form the foundation of the method and will be discussed in more detail later. Following the initial three steps, a collection of 3D coordinates corresponding to points along the cell’s flagellum is obtained. Temporal dynamics are incorporated into the 3D tracking method by iteratively applying the lateral and axial localization modules to subsequent frames within the imaging sequence. The fourth and final module involves converting this collection of 3D coordinates into a quantitative representation of 3D movement. Although we developed and here describe the method specifically for flagellar tracking, it is adaptable for applications involving the 3D movement of entire cells.

**Figure 1.**
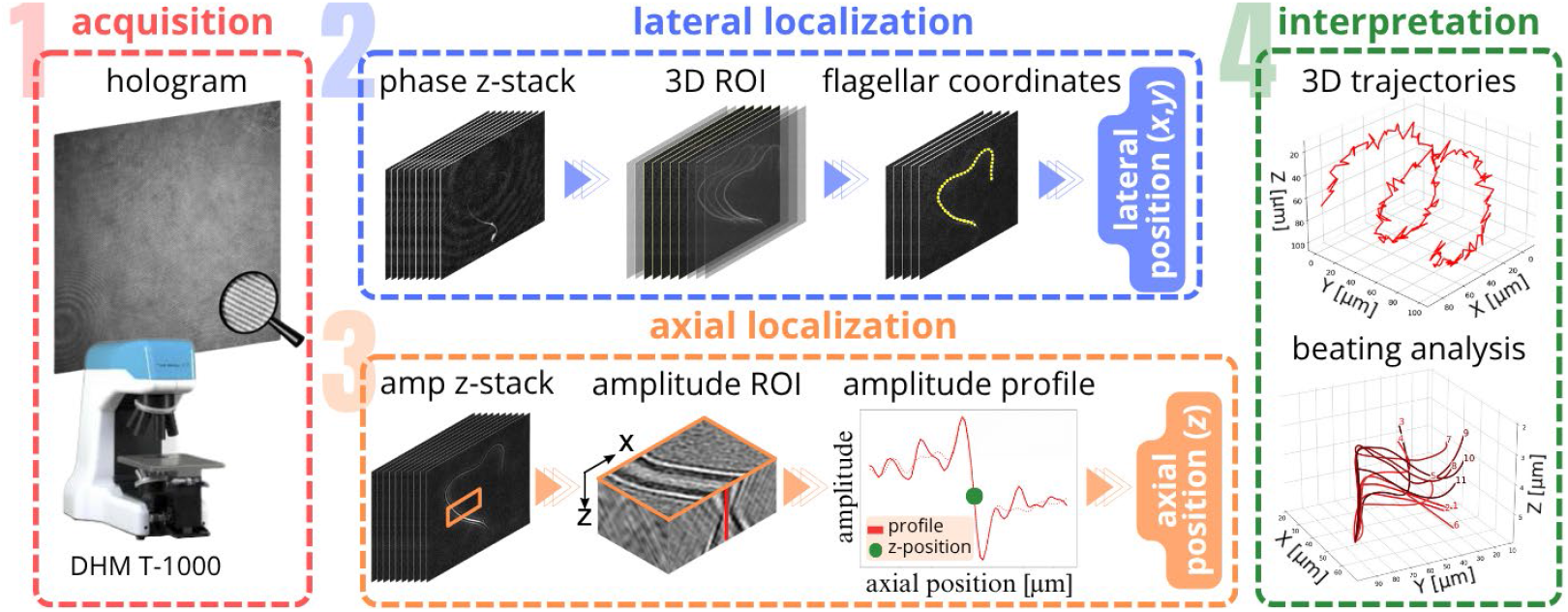
Schematic overview of the holoV3C tracking method using digital holographic microscopy, illustrated for tracking of the mouse sperm cell flagellum. (1) Acquisition: measurement and acquisition of a hologram using a digital holographic microscope. (2) Lateral localization: phase *z*-stack reconstruction, filtering and localization of the 3D phase region of interest (ROI) around the microstructure, and detection of lateral (*x,y*) positions. (3) Axial localization: amplitude *z*-stack reconstruction, processing of the amplitude ROI using (*x,y*) positions from step 2, construction of the amplitude profile along the optical axis and detection of Gouy anomaly. (4) Interpretation: reconstruction of 3D trajectories and flagellum beating analysis.

The modularity of the method is a deliberate design feature aimed at making the approach versatile. This architecture not only promotes scalability but also supports customization of individual modules, allowing it to be modified to meet the specific requirements of diverse experiments. For instance, the lateral localization module can be refined to enhance detection accuracy, accommodating variation among species in flagellar number, arrangement, waveform, and kinematics.

#### 2.1.1. Acquisition

Digital holography provides the foundational data for 3D tracking by enabling precise volumetric imaging. Unlike scanning-based methods, DHM instantaneously captures the entire sampling volume in a single holographic interference pattern, preserving full spatial information within the recorded wavefront. This technique exploits the coherence properties of light by using the interference between two coherent beams: the object beam and the reference beam. As light passes through the specimen, it is collected by a microscope objective to form the object wave. This wave then interferes with the reference wave to generate a hologram, which is recorded by a digital camera. In holoV3C, we employed an off-axis configuration, where the object and reference beams interfere at a small angle. This arrangement facilitates the separation of spatial frequencies [31], streamlining the reconstruction of both amplitude and phase images. Amplitude images are analogous to those observed in transmission light microscopy, whereas phase images represent the product of the cell thickness and the difference in refractive index between the cell and its surrounding medium [32]. The digital reconstruction process of DHM involves emulating the illumination of the hologram using a computationally generated reference wave. This is coupled with numerical adjustments to correct for any wavefront distortions introduced by the optical system, ensuring accurate reconstructions [17, 33].

#### 2.1.2. Lateral localization

Determining the 3D coordinates involves two processes: lateral and axial localization. Following hologram acquisition, the second step consists of accurately identifying the lateral (*x,y*) positions of points on the flagellum. This component fundamentally relies on a 2D tracking framework. Yet, in contrast to 2D tracking which examines a single focal plane per frame, 3D tracking necessitates the analysis of a *z*-stack of images, which complicates the computational process. The principal challenge is the global segmentation of the *z*-stack. When in motion, the flagellum typically spans different heights within the sample volume, so that simply selecting a single plane for image segmentation and subsequent localization of the (*x,y*) coordinates is insufficient. Furthermore, *z*-stack processing methods that are employed in imaging techniques with high specificity, such as fluorescence microscopy, are not directly translatable to holographic imaging due to the inherent noise in the latter.

In holographic imaging, simultaneously capturing images of strongly and weakly scattering objects presents a significant challenge. This issue is particularly evident in flagella tracking, where the strongly scattering cell body can obstruct accurate processing of the weakly scattering flagellum. Furthermore, the appropriate method for accurately determining lateral positions, for a cell or a structure such as a flagellum, depends on various factors, including the morphology of the tracked object, the sampling volume, and the scattering efficiency. Due to these complexities, designing a universal lateral localization pipeline for flagella, which exhibit diverse morphologies and dynamics, remains challenging. In this study, we provide a description of a fully automated lateral localization module specifically designed for the flagella of mouse sperm cells, optimized for their unique morphology and kinematics (Figure 2). In contrast, the processing of *R. americana* flagella required a semi-automated approach due to differences in structural and dynamic properties between the two flagella types. The motion of the flagella of *R. americana* often leads to overlap of flagellar segments, making fully automated segmentation unreliable. To address this, the semi-automated method requires the user to manually define the lateral points of the flagellum in each frame, guiding the algorithm in refining and tracking the flagellum. While both cases adhere to the same general tracking framework, the lateral localization module was adjusted for each type of flagellum to account for its distinct segmentation and processing needs.

**Figure 2.**
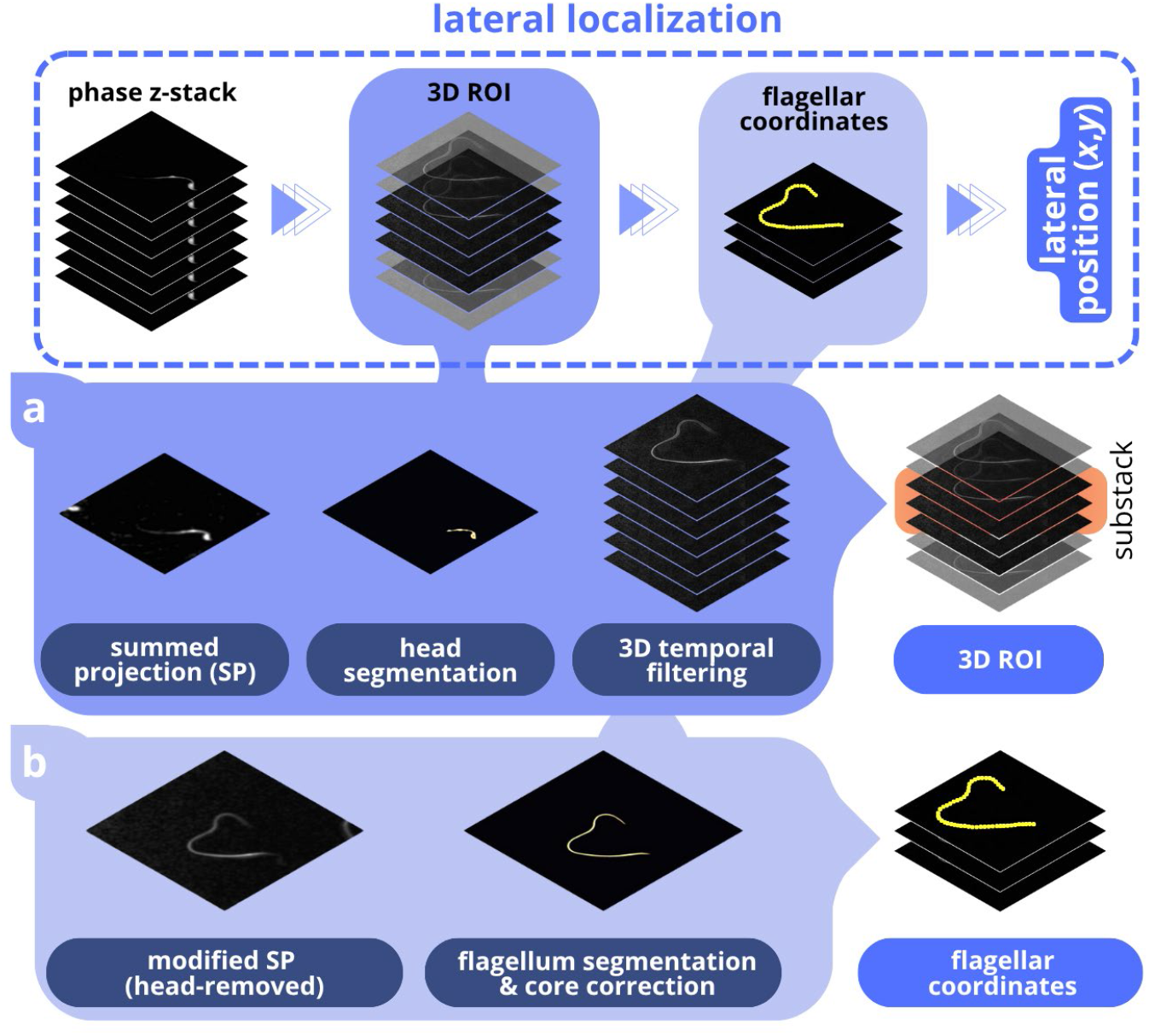
Schematic illustration of the lateral localization module (step 2 in Figure 1) for sperm flagellum analysis. (a) Initial processing of the raw phase *z*-stack uses the Summed Projection (SP) method. Following this, the projection result undergoes segmentation, allowing identification of the sperm head’s lateral (*x,y*) position. These coordinates are integral in defining the focal plane for the sperm head. Subsequently, the *z*-stack undergoes a 3D temporal filtering process aimed at eliminating static background elements, thereby enhancing visibility of the dynamic flagellum. Using the three-dimensional (*x,y,z*) coordinates of the sperm head, the corresponding 3D region of interest (ROI) is established. (b) SP is reapplied to the temporally filtered 3D ROI, facilitating projection of the flagellum’s 3D structure onto a 2D plane. This projected image is then processed through segmentation and flagellum core correction algorithms to refine the representation of the flagellum. The refined segmentation of the flagellar core enables determination of the flagellum’s lateral (*x,y*) positions.

The hologram reconstruction process yields two distinct types of information: amplitude and phase. To accurately determine the lateral coordinates (*x,y*) of the sperm flagellum, we employ the phase *z*-stack, which provides enhanced contrast and better signal-to-noise ratio. The first objective is to define the 3D Region of Interest (ROI) that contains the cell (Figure 2a). The process begins with the initial identification and localization of the cell within the sample volume. To facilitate this, a commonly used technique for reducing the dimensionality of the image *z*-stack is employed, namely, Summed Projection (SP) [34]. Although this method may not be universally suitable due to its potential to reduce the signal-to-noise ratio for certain samples, it remains a reliable technique for the processing of most 3D images. Furthermore, when integrated with additional image processing steps, SP has proven to be effective for holographic images. Head segmentation is then achieved by applying a series of image processing techniques on the summed image. The image is first binarized using Yen’s thresholding method [36]. Next, morphological closing, small object removal, and border cleaning are applied to refine the binary image. From the cleaned image, the medial axis and distance transform are computed, and the coordinates corresponding to the maximum distance are used to localize the lateral position of the sperm cell head. When coupled with the axial localization algorithm described below, this yields the 3D position of the sperm head.

The eukaryotic flagellum poses a significant challenge for image processing due to its slender morphology and rapid dynamics, which often result in diminished contrast and blurring. To address these issues, we implemented 3D temporal filtering to the phase *z*-stack sequence, thereby enhancing the signal-to-noise ratio and improving the visibility of the beating flagellum (Figure 3). This step removes background, artifacts coming from the experimental system, and stationary noise [35]. Additionally, 3D temporal filtering can be employed to compute the 3D (*x,y,z*) coordinates of stationary objects in the sample, aiding in the interpretation of movement by providing contextual information about the surrounding environment. Unlike the conventional temporal filtering algorithm often used in image analysis, 3D temporal filtering is performed across all images in the *z*-stack, not just a single plane. A mean image is calculated for each *z*-stack plane, collectively forming a background *z*-stack for the entire sequence of frames. This background is then subtracted from each *z*-stack, resulting in improved contrast and visibility of objects of interest, including the beating flagellum (Figure 3c).

**Figure 3.**
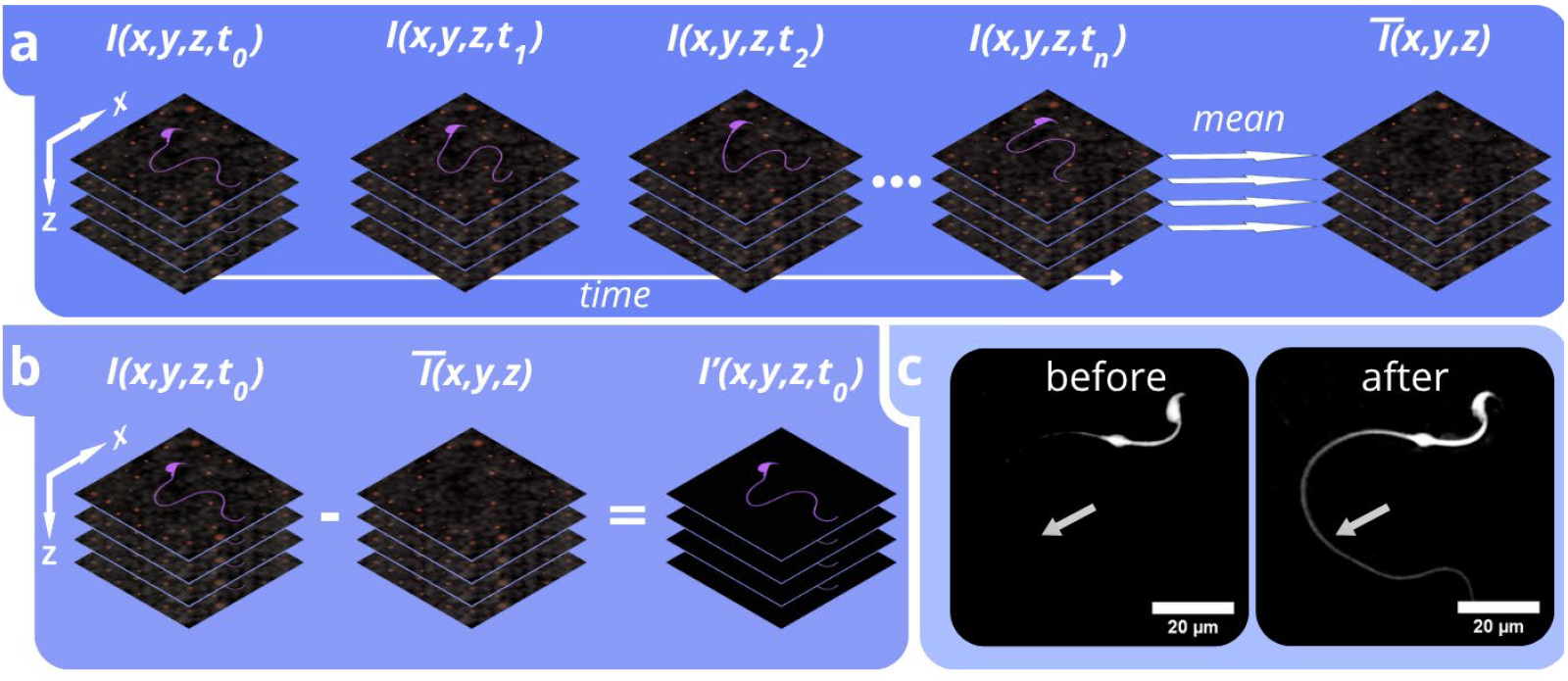
3D temporal filtering of a *z*-stack. (a) Background *z*-stack calculation method. For each corresponding plane across all frames (*t*_0_, *t*_1_, *t*_2_,…,*t*_*n*_), the mean image is computed. These averaged images are then compiled into a new *z*-stack - the background *z*-stack. (b) The second stage of 3D temporal filtering involves subtracting the calculated background *z*-stack from the analyzed stack for each frame. In this way, background and static elements are removed for each plane of the *z*-stack. (c) Illustration of the filtering effect for one *z*-stack plane. The process significantly enhances the visibility of flagella, particularly in segments further from the cell head (indicated by arrows).

By localizing the sperm cell head and midpiece within the sample volume in each frame, we define a 3D ROI in each frame that keeps the center of the head stationary while capturing the region surrounding the tracked flagellum. The lateral dimensions of this ROI can be adjusted as needed to ensure that it contains the entire flagellum. The axial dimensions of this substack depend on the optical system used and the reconstruction parameters. At this point, we have acquired a 3D ROI that has undergone a filtration process to enhance the signal-to-noise ratio. The prepared substack, optimized for clarity and detail through the preceding processing steps, serves as the foundation for the subsequent steps of lateral localization (Figure 2b). Summed Projection (SP) is applied again to compress the 3D ROI into a single 2D image. The prior application of temporal filtration is important because it reduces background noise, resulting in a 2D image where the morphology of the flagellum is sharply defined. This enhanced image is then used for the subsequent step of segmentation.

Segmentation is performed through a sequence of image processing steps to enhance contrast and isolate the flagellum. First, the summed projection image is smoothed using a Gaussian filter, followed by intensity rescaling to optimize contrast. The flagellum is segmented using Yen’s thresholding method [36]. Next, binary dilation and small-object removal are applied as essential morphological operations to refine the structure and eliminate artifacts. The resulting binary image is then skeletonized using Lee’s algorithm [37] to extract the core of the flagellum of one pixel width. Finally, the skeleton points are sorted using a proximity-based approach, ensuring a coherent spatial ordering from the flagellum’s base to its tip. The core is further refined through a correction process that identifies the pixel with the highest intensity within the perpendicular neighborhood of each flagellar point, based on a Maximum Intensity Projection (MIP) [34] image, enhancing positional accuracy. This correction improves lateral localization and reduces the incidence of inaccurately identified points. Through these processing steps, we extract a series of *x,y* coordinates corresponding to points along the flagellum. The number of points varies with the length of the flagellum and the magnification of the objective, as the method attempts to identify every pixel along the flagellum. This effectively represents a 2D tracking framework. By integrating it with the axial localization algorithm, described in the subsequent section, we reconstruct the flagellum’s 3D structure.

#### 2.1.3. Axial localization

Axial localization is a fundamental component in 3D tracking algorithms, permitting the shift from 2D to 3D descriptions of movement. Several existing methods can be used to find the position of a cell along the optical axis. Some require the preparation of reference libraries [10] or calibration processes [11, 13, 14], which can compromise tracking accuracy and prevent them from providing a universal solution across different measurement scenarios. Others quantify image sharpness to determine position [38], but these approaches do not work well for thin and complex structures such as the flagellum. Methods that apply deep learning in quantitative phase imaging, particularly for 3D tracking of cells, are emerging as a promising and innovative approach [39]. However, the efficacy and accuracy of these deep learning methods are contingent on the quality and relevance of the training data. We here introduce a novel method that circumvents the constraints of existing approaches. The fundamental principle of our method lies in simplifying the problem by transforming it from a 3D analysis of the image *z*-stack resulting from holographic reconstruction to a 1D vector analysis of the amplitude profile along the optical axis.

In the lateral localization step, we have identified points (*x,y*) that belong to the flagellum. Using these coordinates, we analyze the amplitude profiles along the optical axis at these positions through the reconstructed image *z*-stack. The processing entails the examination of the amplitude profile to find the location of the Gouy phase anomaly (Figure 4). The theory of phase anomaly states that an additional π phase shift occurs during the wave’s passage through the geometric focus [40]. By identifying the location of this shift, the focal plane can be detected using a single amplitude profile along the optical axis, reducing the analysis of a 2D diffraction pattern to that of a 1D profile and yielding computational savings. This is particularly beneficial given the amount of data generated in the holographic reconstruction, where a single frame can reach a size of up to 540 MB, depending on the reconstruction parameters. Moreover, the method eliminates errors stemming from erroneous choice of the sectioning plane or the presence of noise in the image.

**Figure 4.**
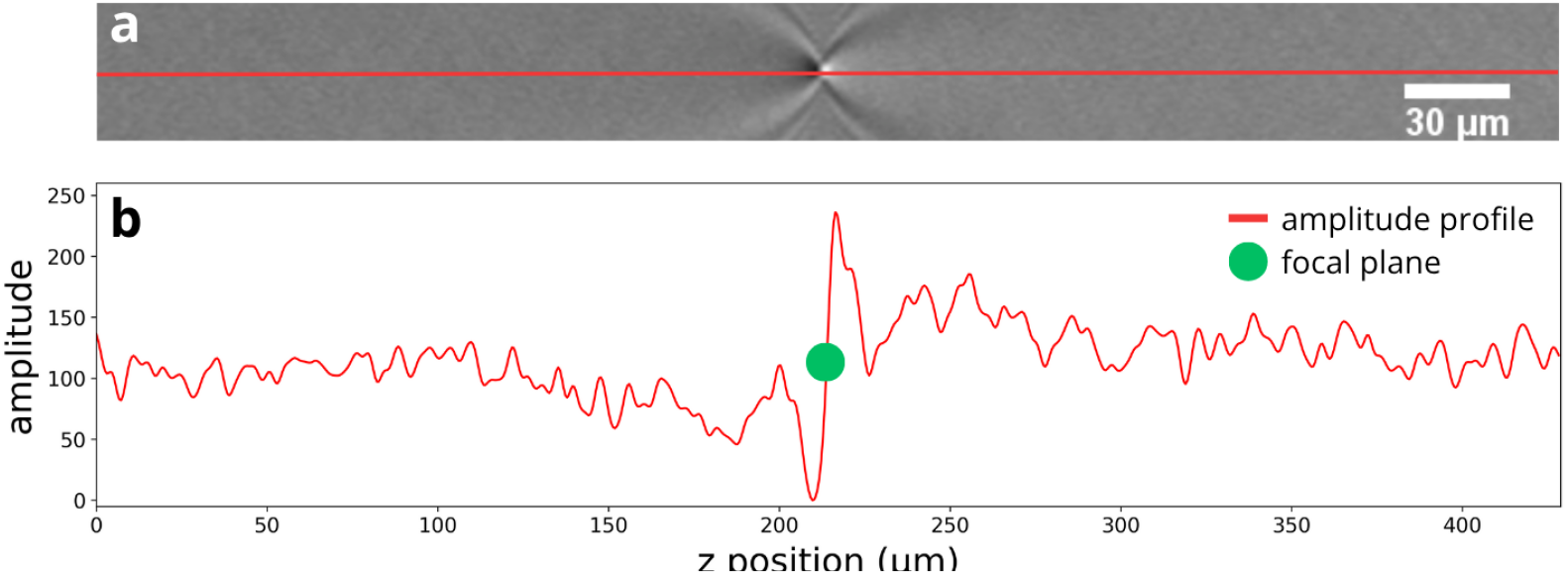
Axial localization (step 3 in Figure 1) ensures precise tracking of samples at extensive depths, maintaining high accuracy. (a) Cross-section of the sample volume along the optical axis, with a red line passing through the center of the point spread function (PSF), marking the profile line used for axial localization. The depth of the reconstructed *z*-stack in the sample plane is 428 µm with a step (distance between *z*-stack planes) of 1 µm. (b) Amplitude profile (red) through the center (corresponding to the *x,y* coordinates determined by lateral localization) of the PSF in panel (a), with the result of the holoV3C algorithm, i.e., the identified *z* position, marked in green.

Further enhancement of the precision of the algorithm is achieved by leveraging the continuous nature of light along the optical axis, allowing for interpolation during the processing of a 1D profile. The method for detecting Gouy’s phase anomaly involves several steps. First, 3D temporal filtering is applied to the amplitude *z*-stack to reduce noise and background information. Next, the amplitude profile is generated by sampling along the optical axis at the cell’s *x,y* position. The profile is then smoothed using a 1D Gaussian filter [41] and interpolated to achieve finer resolution. Finally, the *z*-position is identified as the inflection point near the maximum of the first derivative of the amplitude profile.

Our algorithm enables the precise determination of the axial positions (*z*) based on the lateral (*x,y*) coordinates identified in the preceding step. The approach supports digital refocusing across sample depths extending to millimeters [18]. By analyzing 1D amplitude vectors, this method permits the precise detection of *z*-coordinates over a considerable sampling depth. In the example presented here, the algorithm pinpointed the *z* position within an imaging volume that had a height of 428 μm (Figure 4). This capability effectively harnesses the imaging potential of DHM to satisfy another prerequisite for reconstruction of 3D structures, namely the need for a sufficient sampling volume. The constraints on the dimensions of the imaging volume are primarily determined by the optical parameters of the microscope objective. However, by adjusting holographic reconstruction parameters, it is possible to extend the axial dimension, thereby substantially increasing the accessible imaging volume [42].

#### 2.1.4. Interpretation

The final stage of the holoV3C method transforms the raw 3D points localized in the preceding stages into parameters tailored to the study’s specific objectives. This stage is highly adaptable, allowing one to modify or expand the suite of parameters. Spatial and temporal data are synthesized to enable the quantitative analysis of flagellar beating patterns, allowing the extraction of metrics such as beat frequency, amplitude, and propagation direction. This adaptable framework simplifies the introduction of new parameters to better capture the functional properties of cells or flagella. In this way, the final module ensures broad applicability across different research contexts and permits continuous refinement of the analysis pipeline. Examples of this module, showcasing its application to the analysis of flagellar dynamics, are presented in the following section.

### 2.2. Experimental evaluation

To evaluate the effectiveness of our approach, we analyzed two distinct cell types: mouse spermatozoa and the protist *R. americana*. These cells were selected due to their different morphologies and flagellar beating patterns, which together provide a test of the algorithm’s versatility. Mouse spermatozoa, characterized by a relatively wide (1 µm) and long flagellum (100 µm), may be treated as a benchmark for 3D tracking methods, especially given that most current methods for 3D tracking of flagella focus on sperm flagella. In our experiment, a single sperm cell’s body was naturally adhered to the glass slide. However, this did not affect the lateral localization, as the sperm head’s position was determined for each frame, regardless of its attachment. Our method successfully detected the full length of the mouse sperm flagellum recorded at 10 Hz, yielding approximately 600 points along the flagellum in each *z*-stack (Figure 5). Detection of the full length was confirmed by visual inspection and by the good agreement between our average value of flagellar length (102.9 µm) and that reported in the previous study (96.5 µm) [43].

**Figure 5.**
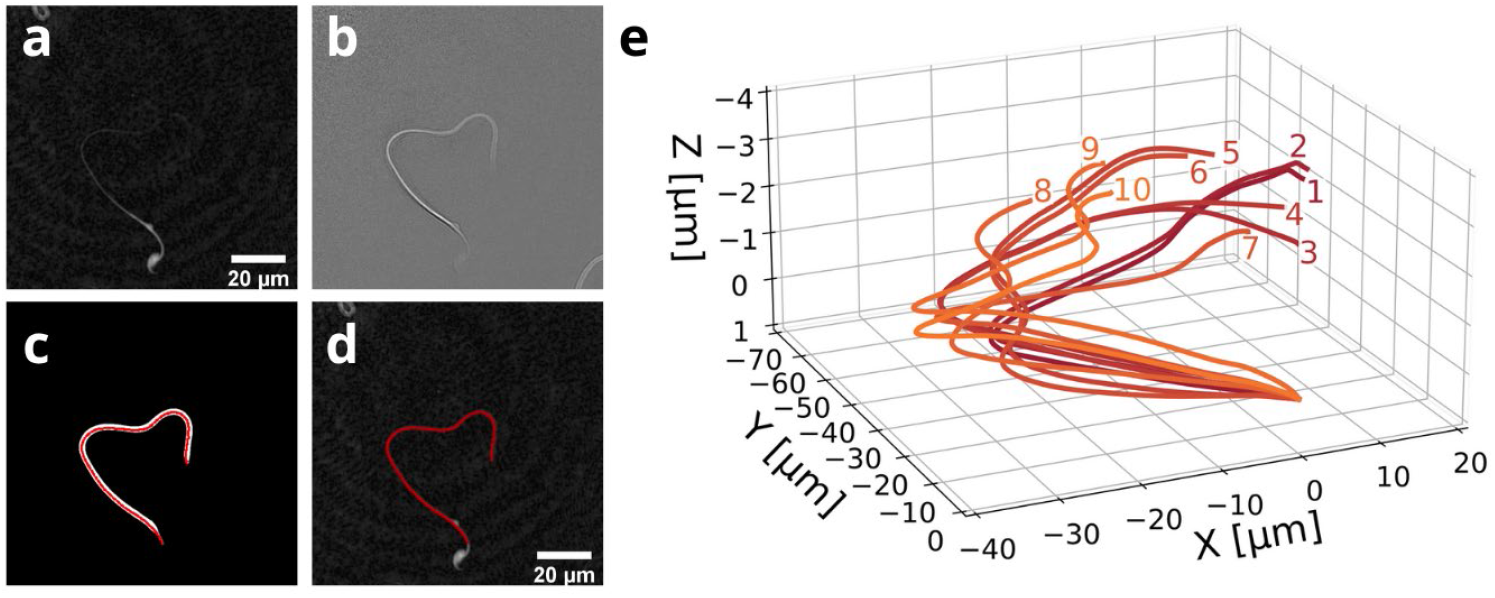
Results from the tracking algorithm reveal the precise movement of the mouse sperm cell flagellum. (a-d) Image processing and segmentation of a sperm flagellum in the *xy* plane. (a) Phase reconstruction on a single *z* plane with visible sperm cell. (b) Effect of 3D temporal filtering. Most of the background, noise, and stationary elements have been removed by this step, improving flagellum visibility. (c) Result of the flagellum core detection algorithm (red). (d) Flagellum core points (red) superimposed on the raw phase image. (e) Flagellum of the same cell plotted as a temporal sequence in 3D space. The illustration shows a 1 s sequence of flagellar movement, with numbers representing ten consecutive frames.

Integrating the lateral and axial localization modules enabled the effective reconstruction of the 3D shape of a mouse sperm cell flagellum and thus its pattern of movement through time (Figure 5e and movie S1). The large number of reconstructed points guarantees a high fidelity in capturing the flagellum’s shape. Thanks to the *O*(*N*) computational complexity of our algorithm, the impact of the number of points and reconstruction parameters on processing speed is linear, keeping the time to process a single frame low. Using a standard PC (3.50 GHz Intel Xeon E5 v4, 24 GB RAM, solid-state drive), our algorithm processed a 32-bit image stack in approximately 2.78 seconds per frame–1.07 seconds for lateral and 1.71 seconds for axial localization. For axial localization, each flagellar point was computed separately, with an average processing time of 0.003 seconds per point. This application validates the applicability and efficacy of the holoV3C method and underscores its capabilities in quantifying the movement dynamics of spermatozoa. Moreover, by incorporating the 3D position of the sperm cell head, determined during the lateral localization stage, our approach provides a tool for the study of motility by integrating 3D tracking of both the head and the flagellar beating pattern.

For our second application, to the flagella of the protist *R. americana*, an organism whose highly non-planar flagellar beating patterns present a critical challenge to existing 3D tracking methods, as discussed earlier. In this experiment, *R. americana* was observed in its feeding stage, attached to the substrate, with its anterior flagellum exhibiting complex 3D beating. These measurements were conducted at 200 fps. Our primary objective was to assess the performance of the axial localization algorithm in the case of a very different morphology and a flagellar width five times smaller than that of mouse sperm cells. The minute size of the flagella, their complex 3D shape, which includes more segments oriented out of plane compared to the mouse sperm cells, and their reduced scattering efficiency together considerably diminish flagella visibility, thereby complicating image *z*-stack processing. Even in the face of these challenges, our method demonstrated its proficiency in capturing the 3D shape of the flagella (Figure 6 and movie S2), confirming its applicability to structures with diameters as small as 200 nm, a size typical for many phagotrophic flagellates [44]. Specifically, we collected approximately 53 points along the flagellum in each *z*-stack, and the measured anterior flagellum length was 17.8 µm. Although direct comparisons are challenging due to the limited existing studies on precise *R. americana* flagellum morphology, these quantitative data highlight our method’s unique capability to resolve such minute and complex flagellar structures. A particular obstacle for 3D tracking was the unique beating pattern of *R. americana*, characterized by numerous instances where parts of the flagella were aligned nearly parallel to the optical axis - a scenario that complicates tracking with many existing methods. Nonetheless, our approach was able to deal with this complication and reconstruct the entire structure, demonstrating its promise as a tool for precisely capturing highly non-planar 3D flagellar kinematics, a key challenge in microbial motility studies [45].

**Figure 6.**
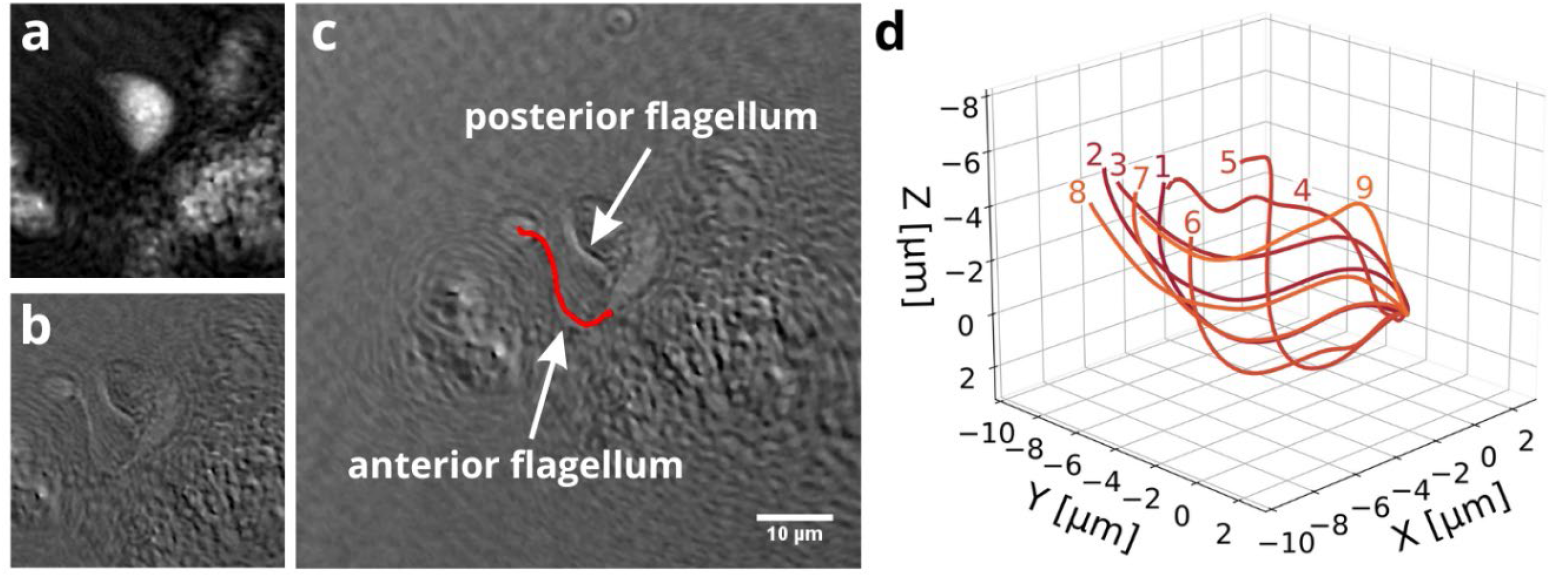
Tracking of the flagella of the protist *R. americana* using the holoV3C algorithm enables high spatiotemporal resolution of the beating pattern. (a) Raw phase reconstruction on a single *z* plane showing strong signal from the body of *R. americana*; the flagella are invisible. (b) Effect of 3D temporal filtering on a single *z* plane. The stationary signal was removed by this processing, which improved the visibility of the flagella. (c) Full field of view phase reconstruction on a single *z* plane. The result of the lateral localization of the anterior flagellum is marked in red. The posterior flagellum is also visible. (d) Flagella of the same cell plotted in 3D space. Numbers represent the consecutive frames of the sequence at 5 ms intervals.

### 2.3. Axial resolution

Axial resolution is a critical yet under-reported parameter for evaluating the performance of 3D tracking methods. It quantifies the precision with which axial positions are calculated, serving as a key indicator of the performance of a 3D tracking method. The ability to define positions along the optical axis is what fundamentally differentiates 3D tracking from 2D tracking. Consequently, accurate reporting of axial resolution is essential to characterize 3D tracking techniques and to enable meaningful comparisons between methods. However, axial resolution is influenced not only by the tracking method itself but also by other factors such as sample morphology, scattering properties, or immobilization. These factors are often overlooked but can significantly impact the axial resolution. Building on existing strategies for determining axial resolution in 3D tracking methods [46, 47], we define axial resolution as the root mean square error (RMSE) of axial positions. For each tracked point, we determine its *z*- coordinate and compare it to a corresponding reference position (described below). The residual error, given by the difference between the measured and reference *z*-coordinates, is then used to compute the RMSE. In this section, we investigate how a single 3D tracking algorithm can produce different axial resolutions when applied to different tracking subjects, namely polystyrene particles and mouse sperm flagella. This comparison illustrates the interplay between algorithm performance and sample-specific characteristics, emphasizing the need for context-dependent assessment of the axial resolution to ensure accurate and reliable evaluation.

Small, spherical particles are the most commonly used morphology for the estimation of axial resolution [10, 13, 30, 46, 47], making them an ideal starting point for our analysis. We conducted measurements on randomly dispersed polystyrene particles (diameter 1.05 µm, SD = 0.03 µm, microParticles) suspended in deionized water, at different sampling times (5 frames, Δ*t* = 94 ms). For each frame, 30 particles were analyzed, and for each particle, a total of five lateral points were determined within the boundary of the particle: one central point and four peripheral points, arranged in a configuration known as the Von Neumann neighborhood [48]. These points, across the five sampling times, resulted in a total of 750 data points. The selection of the central point was conducted manually, a decision intended to avoid algorithmic bias, coupled with the consideration that lateral resolution does not constitute the primary focus of this analysis. For each designated (*x,y*) point, its position along the optical axis (*z*) was determined using the axial localization module. These calculated *z*-positions were then averaged for each particle, providing the reference values used for residual calculations. The mean positions were subsequently validated through visual inspection. The decision to opt for the mean position, rather than a visually determined reference position, was made not only to minimize potential errors associated with the manual identification of each particle’s focus position, but also to reduce errors related to the imaging system itself. After the residual calculation for each particle, based on the local average, an outlier elimination process was employed, leveraging the Interquartile Range (IQR) method [49] to ensure the integrity of the data set. Our method achieved an axial resolution, calculated as the RMSE, of 0.053 µm for the polystyrene particles (Table 1).

**Table 1.** Performance in axial localization of the 3D tracking method.

| | Root mean square error<br>($\mu\text{m}$) | Mean bias error<br>($\mu\text{m}$) | Maximum error<br>($\mu\text{m}$) |
| --- | --- | --- | --- |
| Flagellum | 0.247 | -0.016 | 0.676 |
| Particles | 0.053 | 0.003 | 0.131 |

Motivated by observed disparities in *z*-precision between latex bead measurements and 3D flagellar reconstructions [13], we also assessed the resolution of our method when applied to flagella. To achieve this, we measured the flagellum of a mouse sperm at 35 sampling times (Δ*t* = 94 ms). At each sampling time, 500 flagellum points were identified, resulting in a total of 17500 flagella points. The lateral (*x,y*) positions were determined manually, thereby circumventing any potential inaccuracies attributable to the lateral localization algorithm. Given the difficulty of visually determining the *z*-position of individual flagellum points, reference positions were obtained by smoothing the data in the *z*-dimension. A Savitzky-Golay filter [50] (window size 95, polynomial order 3) was applied along the *z*-axis to obtain a reference flagellum shape. This smoothed flagellum served as the basis for calculating residuals for the 17500 points, after which outlier points were removed using the Interquartile Range (IQR) method [49]. We then computed the root mean square error, and for further comparison of performance between the analysis of flagella and particles, calculated the mean bias error [49] and the maximum localization error for both datasets (Table 1).

Our method achieved an axial resolution (Root Mean Square Error, or RMSE) of 0.053 μm for polystyrene particles and 0.247 μm for the sperm cell flagellum, demonstrating its precision in 3D tracking applications. For both samples, the tracking method exhibited a low value of Mean Bias Error (MBE). These minimal MBE values indicate negligible systematic bias, which emphasizes the method’s adaptability and reliability across various sample types. Moreover, the maximum error values were low, at 0.676 μm for the sperm flagellum and 0.131 μm for particles. These low values indicate that the axial localization method maintains minimal deviations from true positions. These metrics collectively underscore the method’s broad applicability and high precision. Despite the low values of the MBE and maximum error, we observed a considerable difference in the Root Mean Square Error (RMSE) between the two sample types. The observed differences between the RMSE results for the stationary particles and sperm flagella show that the same tracking method behaves differently for different test objects. Sample characteristics as well as measurement system parameters such as image pixel size or uniformity of the illumination have a significant effect on the axial resolution [51]. Inaccurate estimation of axial resolution can lead to misinterpretation of biological dynamics, error propagation in quantitative models, and ultimately misleading comparisons across studies. Recognizing the challenges in developing a universal procedure for determining the axial resolution of tracking methods, we highlight the importance of addressing this issue. Establishing community-agreed protocols would facilitate the evaluation and comparison of algorithms, and clarify the range of their applicability. Furthermore, the creation of benchmark datasets tailored to different tracking scenarios would serve as a valuable resource for validating and refining axial localization techniques.

## 3. Discussion

Understanding the complex, often highly non-planar, 3D structure and movement of flagella is important for unraveling the mechanisms underlying cellular motility across diverse applications [4]. However, this morphological diversity of flagella, particularly with significant out-of-plane components, poses technical and computational challenges [52]. To address these challenges, a method must combine high spatial and temporal resolution, the ability to sample large volumes, and computational efficiency to ensure robust and reliable performance across diverse biological contexts.

In this study, we have presented a method that uses digital holographic microscopy to overcome this challenge. Our method demonstrates high axial resolution, achieving 0.247 μm for beating mouse sperm flagella and down to 0.053 μm for polystyrene particles, along with precise 3D localization at the single-pixel level across large sample depths. This capability is a critical feature for accurately resolving the full, non-planar 3D flagellar kinematics. Furthermore, the system achieves measurements at high temporal resolution, with a capability of up to 200 frames per second. This frame rate does not represent an inherent limitation of the method, but rather the upper boundary of the imaging camera we used. The existence of commercial DHM setups capable of capturing images at speeds of up to 100,000 frames per second illustrates the scalability and potential of this approach for imaging at even higher speed in the future. The computational efficiency of our algorithm provides a strong foundation for easy implementation in systems equipped with cameras offering higher temporal resolution, enabling further progress in tracking dynamic behaviors such as 3D flagellar beating with enhanced temporal detail.

Our approach demonstrates a significant improvement in axial resolution, calculated as the root mean square error (Table 1), compared to existing 3D tracking methods. The axial resolution achieved by existing methods varies widely depending on the imaging technique. Light microscopy with piezo-driven objectives provides an axial resolution of 3.2 µm, while multifocal imaging has achieved resolutions of 0.15 µm [13] and 0.12 µm [11]. Other DHM methods have demonstrated resolutions of 0.15 µm [29, 30]. However, many techniques do not explicitly report axial resolution, making direct comparisons difficult and underscoring the need for standardized benchmarks [6, 12, 26–28]. In contrast, our holoV3C algorithm achieves an axial resolution of 0.05 µm (Table 1), offering a threefold improvement over multifocal and DHM methods and surpassing piezo-driven light microscopy by more than an order of magnitude. We also found that the axial resolution in 3D tracking is related to the properties of the object being tracked (Table 1). This dependency is likely to be driven by multiple factors, encompassing the geometrical shape, volumetric dimensions, scattering efficiency, material composition, and the motility characteristics of the object. Accordingly, we propose that estimates of the resolution should be conducted with materials and conditions that are specific to the application of interest to ensure accurate quantification.

A strength of digital holographic microscopy, enhanced by the proposed algorithm, is the ability to achieve 3D tracking in large sampling volumes. We were able to precisely reconstruct the 3D dynamics of a sperm flagellum over a depth of 428 μm. This represents a 26-fold increase compared to a previously proposed DHM method [26] and a 21-fold increase compared to multifocal imaging [13]. Expanding the reconstruction range along the optical axis enables the capture of 3D flagellar beating patterns that are characterized by significant vertical components, as observed in the protist *R. americana*. This expanded range also enables the measurement of organisms located at different depths within a sample, thereby increasing the throughput of the method.

Our method simultaneously integrates high spatial and temporal resolution with computational efficiency, a combination that is important for broad applicability in biological imaging. Several existing approaches depend on complex computational techniques to achieve high axial resolution, a factor that limits processing speed and scalability. Our algorithm achieves *O*(*N*) computational complexity, offering a significant advantage over alternative methods. Approaches employing holographic deconvolution [19] or fitting Airy ring patterns [20] typically exhibit *O*(*N*log*N*) complexity, leading to higher computational costs. Some methods even reach *O*(*N*²) complexity [21, 22], imposing substantial computational demands and further limiting their practical applicability. These differences in computational complexity are particularly important in the context of flagellar tracking, where a large number of points, approximately 600 per time point in our sperm cell flagellum application, are used to accurately resolve the flagellum’s shape. Consequently, an *O*(*N*) algorithm will have a ninefold improvement in efficiency compared to an *O*(*N*log *N*) algorithm, allowing for more streamlined processing of larger datasets and enhanced scalability. In future implementations, lateral and axial localization steps could be offloaded to a graphics processing unit (GPU), achieving significant processing time improvement. Importantly, because image acquisition and localization are separated stages (Figure 1), even current processing times do not limit the acquisition rate, which remains the principal bottleneck in tracking rapid dynamic movements.

Applications of our method to a sperm flagellum and, crucially, to the protist *R. americana* have demonstrated its effectiveness and robustness in accurately quantifying and measuring the full 3D shape and highly non-planar kinematics of flagella over time. By combining high temporal and spatial resolution with computational efficiency, this method provides a robust framework for the comprehensive characterization of complex, non-planar flagellar dynamics that often cannot be fully resolved by conventional 2D or other 3D tracking approaches. This makes it promising for a broad range of applications in microbiology, notwithstanding the need to adapt certain aspects (e.g., the quantification of axial resolution) to the specific context. The ability to examine highly non-planar 3D flagellar dynamics can enhance our understanding of cellular locomotion and its biological and ecological functions. Beyond the analysis of flagella, the method has wider potential applications, including 3D tracking of bacterial motility. Realizing this potential would require adapting the lateral localization step of the algorithm to accurately identify and track the centroids of microbial cells, thereby extending its use to other aspects of the dynamics of microbial movement.

## 4. Methods

### 4.1. Ethical statement

All experimental procedures involving animals complied with international ethical standards. Animal research was approved by the École Polytechnique Fédérale de Lausanne (EPFL) Animal Research Ethics Committee under license VD3290.1. All protocols were performed in accordance with relevant guidelines and regulations.

### 4.2. Sample preparation

Mouse spermatozoa samples were kindly provided by Sonia Verp (EPFL). An 11-week-old B6.CD45.1 male mouse was euthanized under approved protocol (VD3290.1) by CO2 inhalation followed by cervical dislocation. Immediately post-mortem, the two cauda epididymides were excised and placed on sterile filter paper. Under microscopic observation, all visible adipose tissue and blood were removed. Using fine forceps, 5–6 incisions were made in each epididymis to facilitate the release of spermatozoa. Epididymides were transferred into individual Eppendorf tube containing preincubation medium (CARD FERTIUP, Cosmo Bio LTD). The tube was incubated to allow the sperm to swim up into the medium. Following incubation, the sperm suspension was collected from the surface using a micropipette, thereby minimizing tissue debris and obtaining the most motile spermatozoa.

Samples of the aerobic freshwater species *Reclinomonas americana* (isolate ATCC50394) were kindly provided by Alastair Simpson (Dalhousie University). Cultures were maintained in the dark at 18°C in Milli-Q^®^ water supplemented with 0.3% Miller’s LB broth media.

### 4.3. DHM imaging

The measuring system used in this study is a transmission digital holographic microscope, DHM™ T-1000 (Lyncée Tec SA, Switzerland), in an off-axis configuration. The setup includes a 666 nm laser light source with illumination delivered through an optical fiber and subsequently collimated, a 40× (NA = 0.75, HCX PL FLUOTAR, Leica) and a 63× (NA = 1.3, HC PL FLUOTAR, Leica) microscope objectives, and a camera (acA1920-155um, Basler AG, Germany) with 5.86 µm/pixel. This setup enabled the acquisition of holograms at a frame rate of 200 frames per second (fps) for the full field of view (1024 × 1024 pixels). Samples were observed in a custom-built, 0.5 mm–high observation chamber, assembled by sealing a microscope slide with a coverslip using rubber spacers.

### 4.4. Algorithm design and implementation

The holoV3C method is structured into four sequential core modules: acquisition, lateral localization, axial localization, and interpretation (Figure 1). Holographic reconstruction of raw hologram data, including phase unwrapping and wavefront propagation, was performed using the Koala^®^ software by Lyncée Tec (version 8.6). The core localization algorithm, including the lateral (*x,y*) and axial (*z*) localization stages, was custom-developed in Python (version 3.11) utilizing standard scientific libraries including NumPy (version 1.26.4), SciPy (version 1.13.1), and scikit-image (version 0.23.2). The integration of these *x,y*, and *z* coordinates for each frame allows for the 3D flagellar reconstruction. Temporal dynamics are then captured by iteratively applying these localization modules to subsequent frames within the imaging sequence, thereby generating a time-resolved 3D representation of the flagellar movement.

### 4.5. Statistical analysis

We quantified axial resolution by computing the root mean square error (RMSE) between measured and reference *z*-positions. For polystyrene particles, reference axial positions were determined by averaging calculated *z*-coordinates across five lateral points within each particle, recorded over five frames (totaling 750 points), and subsequently verified by visual inspection. For mouse sperm flagella, axial reference positions were established by smoothing the raw axial position data using a Savitzky-Golay filter (window size 95, polynomial order 3), applied to the 17,500 measured flagellar points. Outliers were systematically removed using the Interquartile Range (IQR) method. In addition to RMSE, Mean Bias Error (MBE) and maximum error were computed for both datasets to assess systematic deviations and the maximal precision bounds of the method. Statistical analyses were performed in Python (version 3.11) utilizing standard libraries including NumPy (version 1.26.4) and SciPy (version 1.13.1).

## 5. Data availability

The data that support the findings of this study are publicly available on Zenodo at https://doi.org/10.5281/zenodo.13842914

## Acknowledgments

We thank Sonia Verp and Alexandre Widmer (Center of PhenoGenomics, EPFL) for their expertise and for providing mouse spermatozoa samples; Alastair Simpson (Dalhousie University) for providing *Reclinomonas americana* samples; our colleagues at the Stocker lab and the PHYMOT fellows for their feedback that propelled this project forward; and the Lyncée Tec team for their guidance on holographic microscopy and for sparking multiple stimulating discussions. Finally, we thank Elzbieta Sliwerska for her support with logistics and experimental operations, and Russell Naisbit for his valuable editorial input.

## 8. Funding

We gratefully acknowledge financial support from the following sources: the European Union’s Horizon 2020 research and innovation program under Marie Skłodowska-Curie (Grant No. 955910) to P.N.; the Swiss National Science Foundation Ambizione (Grant No. PZ00P2_202188) to J.S.; the Gordon and Betty Moore Foundation Symbiosis in Aquatic Systems Initiative Investigator Award (GBMF 9197); the Simons Foundation’s Principles of Microbial Ecosystems collaboration (Grant 542395FY22); the Swiss National Science Foundation (Grant 205321_207488); Swiss National Science Foundation Sinergia (Grant CRSII5-186422); the Swiss National Science Foundation National Centre of Competence in Research Microbiomes (Grant 51NF40_180575) to R.S.; and a research grant from the Human Frontier Science Program (RGP014/2024) to T.K.

### Contributions

P.N., J.S., Y.E., T.K., and R.S. conceptualized and designed the study. P.N., C.M.P., and F.M. conducted the experimental work. T.C. developed the *z*-stack reconstruction method. P.N. developed the 3D tracking method, performed data analysis, and created visualizations. P.N., J.S., Y.E., and R.S. drafted the manuscript, with input from all authors. All authors reviewed and edited the manuscript before submission.

