## supplementary_materials for "Precise 3D Tracking of Highly Non-planar Eukaryotic Flagellar Beating Patterns using Digital Holographic Microscopy"

Patryk Nienaltowski *et al.*

**This PDF file includes:**

Legend for movies S1 to S2
Legend for data S1 to S6

**Other Supplementary Materials for this manuscript include the following:**

Movies S1 to S2
Data S1 to S6

**Movie S1.**

Three-dimensional tracking of the mouse spermatozoa flagellum. The left clip shows a two-dimensional phase reconstruction on a single *z*-plane after 3D temporal filtering, with lateral localization of the anterior flagellum highlighted in red. Center and right clips show the flagellum's 3D beating pattern at distinct elevation and azimuth angles to illustrate its full kinematics. Video is slowed 2× to enhance visualization of the flagellar beat.

**Movie S2.**

Three-dimensional tracking of the *R. americana* flagellum. The left clip shows a two-dimensional phase reconstruction on a single *z*-plane after 3D temporal filtering, with lateral localization of the anterior flagellum highlighted in red. Center and right clips show the flagellum's 3D beating pattern at distinct elevation and azimuth angles to illustrate its full kinematics. Video is slowed 40× to enhance visualization of the flagellar beat.

**Data S1. (separate file)**

Results of 3D tracking of mouse spermatozoa flagellum. Processed (*x*,*y*,*z*) coordinates for each frame of the analyzed sequence are provided in individual worksheets, named by frame number. The "metadata" worksheet details the experimental settings. The "all_data" worksheet concatenates the full time series of 3D coordinates for the analyzed sequence.

**Data S2. (separate file)**

Results of 3D tracking of *R. americana* flagellum. Processed (*x*,*y*,*z*) coordinates for each frame of the analyzed sequence are provided in individual worksheets, named by frame number. The "metadata" worksheet details the experimental settings. The "all_data" worksheet concatenates the full time series of 3D coordinates for the analyzed sequence.

**Data S3. (separate file)**

Processing time logs for lateral and axial localization in the mouse spermatozoa flagellum 3D tracking pipeline. For each frame, the elapsed times (s) of each core stage of the tracking pipeline are recorded. The worksheet summarizes average and standard deviation for each stage across the full sequence. The "metadata" worksheet details the experimental settings.

**Data S4. (separate file)**

Three-dimensional coordinates of the mouse spermatozoa flagellum used for the axial resolution analysis. Processed (*x*,*y*,*z*) coordinates for each frame of the analyzed sequence are provided in individual worksheets, named by frame number. The "metadata" worksheet details the experimental settings.

**Data S5. (separate file)**

Three-dimensional coordinates of the dispersed polystyrene 1.05-μm research particles used for the axial resolution analysis. Processed (*x*,*y*,*z*) coordinates for each frame of the analyzed sequence are provided in individual worksheets, named by frame number. The "metadata" worksheet details the experimental settings.

**Data S6. (separate file)**

Axial resolution metrics. Consolidated performance table summarizing root-mean-square error (RMSE), mean bias error (MBE), and maximum error in axial localization for both test objects - mouse spermatozoa flagellum (Data S4) and polystyrene particles (Data S5). All metrics are reported in micrometers. The “metadata” worksheet details the experimental settings.
